# High- to low-level decoding does not generally improve perceptual performance

**DOI:** 10.1101/240390

**Authors:** Long Luu, Cheng Qiu, Alan A. Stocker

## Abstract

Ding et al. (1) recently proposed that the brain automatically encodes high-level, relative stimulus information (i.e. the ordinal relation between two lines), which it then uses to constrain the decoding of low-level, absolute stimulus features (i.e. when recalling the actual lines orientation). This is an interesting idea that is in line with the self-consistent Bayesian observer model (2, 3) and may have important implications for understanding how the brain processes sensory information. However, the notion suggested in Ding et al. (1) that the brain uses this decoding strategy because it improves perceptual performance is misleading. Here we clarify the decoding model and compare its perceptual performance under various noise and signal conditions.

---

Orientations of the 2-line stimulus used in Ding et al. (1) (Fig.1a, black asterisk; 50° and 53° orientation) are encoded as two noisy, sensory measurements (Fig.1a, red) that can be further corrupted by memory noise (Fig.1a, blue). A low- to high-level decoding strategy predicts estimates that are identical to the memory samples (assuming a flat prior for simplicity) and are distributed according to the combined sensory and memory noise (Fig.1a, right column). In contrast, a high- to low-level decoder assumes that the line estimates are conditioned on the relative order of the two lines, which predicts the characteristic repulsive bias pattern in the estimates. Importantly, if ordinal information is based on the memory samples, then the low- to high-level and the high- to low-level decoder are identical with regard to the ordinal information contained in the estimates (ordinal performance 68%); that is, a high- to low-level decoding strategy *per se* does not improve ordinal discrimination because ordinal performance is strictly limited by the available stimulus information (Signal Detection Theory). Only if the high- to low-level decoder has access to ordinal information contained in the less corrupted sensory measurements, ordinal performance is higher because the ordinal information contained in the sensory signals is preserved in the estimates.

**Figure 1:**
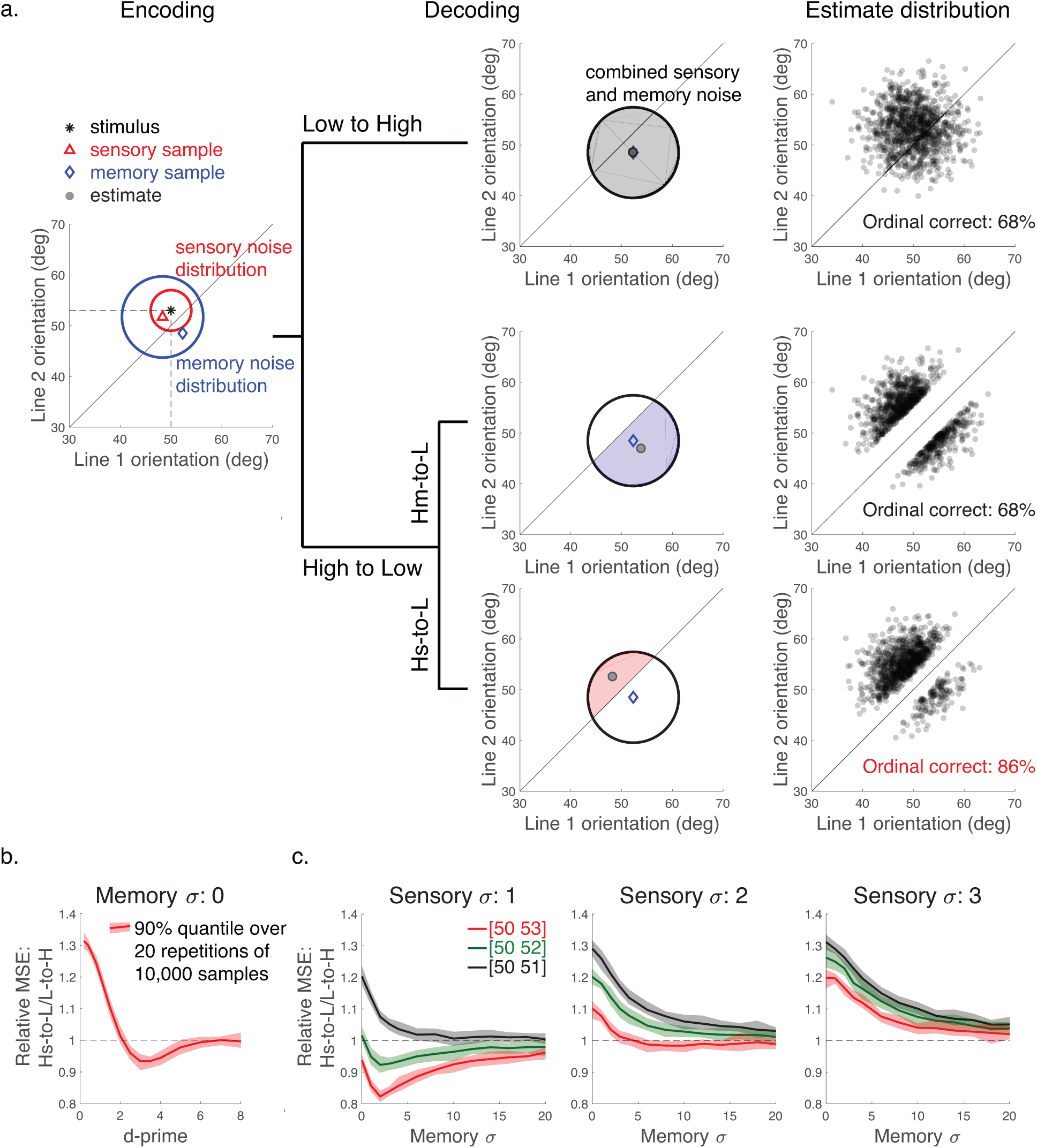
Decoding performance. **(a)** Line orientations are encoded according to sensory and memory noise. Low- to high-level decoding results in orientation estimates that are distributed according to the combined noise. High- to low-level decoding results in characteristic bi-modal distributions. If conditioning is based on memory samples (Hm-to-L) then ordinal information in the estimates is the same as for the low- to high-level decoder. Only if conditioning is based on the less noisy sensory samples (Hs-to-L), ordinal performance improves. **(b)** Estimation accuracy (relative mean-squared-error (MSE)) as function of d-prime. **(c)** Relative MSE for different noise conditions and three pairs of line orientations.

Ultimately, however, subjects in Ding et al. (1) were asked to provide accurate orientation estimates. We computed the relative estimation accuracy between high-to low-level and low-to high-level decoding in terms of mean squared-error for different sensory and memory noise levels. If both decoders have the same sensory information (no memory noise) then high-to low-level decoding only performs better in a narrow regime of intermediate d-prime values (Fig. 1b). When adding memory noise and giving high-to low-level decoding the advantage of having more accurate ordinal information, it also rarely outperforms the low-to high-level decoder in particular for larger sensory noise (Fig. 1c). Therefore, the implication by Ding et al. (1) that the high-to low-level decoding strategy generally improves the reliability of the decoded orientations is incorrect.

In conclusion, Ding et al. (1) provided a great example of self-consistent/high-to low-level inference. However, the benefits of this decoding strategy have been overstated; why and under what conditions the brain adopts such a strategy remains an open (yet very interesting!) question.

## References

(1) Ding S, et al. (2017) Visual perception as retrospective Bayesian decoding from high-to low-level features. Proc. National Academies of Sciences U.S.A 114 (43) E9115–E9124

(2) Stocker AA, Simoncelli EP (2007) A Bayesian model of conditioned perception. Adv. In Neural Information Processing Systems (NIPS) Vol.20: 1409–1416

(3) Long L, and Stocker AA (2016) Choice-induced biases in perception. bioRxiv 043224

